# A risk assessment framework for seed degeneration: Informing an integrated seed health strategy for vegetatively-propagated crops

**DOI:** 10.1101/105361

**Authors:** S. Thomas-Sharma, J. Andrade-Piedra, M. Carvajal Yepes, J. F. Hernandez Nopsa, M. J. Jeger, R. A. C. Jones, P. Kromann, J. P. Legg, J. Yuen, G. A. Forbes, K. A. Garrett

**Affiliations:** Department of Plant Pathology, Kansas State University KS, USA; Department of Plant Pathology, University of Wisconsin-Madison, WI, USA; International Potato Center, Lima, Peru; International Center for Tropical Agriculture, Cali, Colombia; Institute for Sustainable Food Systems, Plant Pathology 12 Department, and Emerging Pathogens Institute, University of Florida, Gainesville, FL, USA; Centre for Environmental Policy, Imperial College London, UK; Institute of Agriculture, University of Western Australia, Crawley, Australia; International Potato Center Quito, Ecuador; International Institute of Tropical Agriculture,Dar es Salaam, Tanzania; Department of Forest Mycology and Plant Pathology, Swedish University of 18 Agricultural SciencesUppsala, Sweden; International Potato Center Kunming, China

**Author notes:** Corresponding authors: K. A. Garrett and S. Thomas-Sharma 22 E-mail addresses and.

**Keywords:** banana, cassava, environmental heterogeneity, positive selection, potato, root crops, seed degeneration, seed health, simulation models, sweetpotato, tuber crops, vegetative propagation, yam.

## Abstract

Pathogen build-up in vegetative planting material, termed seed degeneration, is a major problem in many low-income countries. When smallholder farmers use seed produced on-farm or acquired outside certified programs, it is often infected. We introduce a risk assessment framework for seed degeneration, evaluating the relative performance of individual and combined components of an integrated seed health strategy. The frequency distribution of management performance outcomes was evaluated for models incorporating biological and environmental heterogeneity, with the following results. (1) On-farm seed selection can perform as well as certified seed, if the rate of success in selecting healthy plants for seed production is high; (2) When choosing among within-season management strategies, external inoculum can determine the relative usefulness of ‘incidence-altering management’ (affecting the proportion of diseased plants/seeds) and rate-altering management (affecting the rate of disease transmission in the field); (3) Under severe disease scenarios, where it is difficult to implement management components at high levels of effectiveness, combining management components can produce synergistic benefits and keep seed degeneration below a threshold; (4) Combining management components can also close the yield gap between average and worst-case scenarios. We also illustrate the potential for expert elicitation to provide parameter estimates when data are unavailable.

## INTRODUCTION

In vegetatively-propagated crops, pathogens tend to accumulate if planting material is drawn from within a crop population over multiple generations, resulting in significant quality and yield losses. This problem, termed seed degeneration (where ‘seed’ refers to vegetative planting material), occurs commonly when certified, disease-free planting material is scarce and/or expensive, as is the case in many low-income countries (Gibson and Kreuze 2014; Thomas‐Sharma et al. 2016) and for some specialty crops (Gergerich et al. 2015). An integrated seed health strategy (Thomas‐Sharma et al. 2016) is needed to address seed degeneration, drawing on management components that are currently available to farmers, or can be made available in the near future. We present a risk assessment framework for seed degeneration in vegetatively-propagated crops, designed to support the development of strategies for integrating management components.

Seed degeneration is affected by many biophysical factors such as the susceptibility of a variety, the abundance of alternative hosts (e.g., weeds), the roles and efficiencies of vectors, regional inoculum availability, and the conduciveness of weather for disease development and spread. Processes such as reversion, where seed obtained from infected mother plants is healthy, can reduce seed degeneration in sweetpotato (Gibson et al. 2014), potato (Bertschinger 1992), and cassava (Fargette et al. 1996; Gibson and OtimNape 1997). The etiology of seed degeneration is often specific to a crop and geographical region. Cassava mosaic geminiviruses (CMGs) and Cassava brown streak viruses (CBSVs) are major causes of degeneration in East Africa (Legg et al. 2015), while viruses associated with cassava frogskin disease are the main causes of degeneration in South America (Carvajal-Yepes et al. 2014). For potato, viruses are a major cause of seed degeneration around the world (Thomas‐Sharma et al. 2016), while latenttuber infections of the bacterial wilt pathogen, *Ralstonia solanacearum*, are a major problem in tropical and subtropical countries (Mwangi et al. 2008), and the fungal pathogen *Rhizoctonia solani* is a problem at high altitudes in the Andes (Fankhauser 2000).

The use of seed certified to be disease-free or to have high health status (hereafter referred to as certified seed) is often recommended as the primary management strategy to counter on-farm seed degeneration (Frost et al. 2013; Gergerich et al. 2015; Thomas‐Sharma et al. 2016). Examples of such formally regulated systems include the US National Clean Plant Network (www.nationalcleanplantnetwork.org), which supplies seed material for many fruit crops, the Wisconsin Seed Potato Improvement Association (www.potatoseed.org), which supplies seed potato in Wisconsin, USA, and the Netherlands General Inspection Service for Agricultural Seeds and Seed Potatoes (www.nak.nl), which certifies seed potatoes from the Netherlands for global export. However, specialized programs designed to produce healthy seed are rarely used by smallholder farmers in low-income countries (Thiele 1999; Thomas‐Sharma et al. 2016). In many low-income countries, 80-95% of seed is routinely obtained from informal seed sources with poor or unknown seed health status (Mallowa et al. 2006; McGuire and Sperling 2016; Thomas‐Sharma et al. 2016). Thus, one focus for the application of this new risk assessment framework is the context of low-income countries, especially food-security crops such as banana and plantain, cassava, potato, sweetpotato, and yam.

Evaluating management components, and the potential synergies from combining them, is a key part of a risk assessment framework for seed degeneration. Synergies can be evaluated in terms of reductions in disease, or increases in yield, that are greater when management components are combined compared to the sum of the individual component effects. Seed degeneration can be managed by limiting epidemics in the field, using components such as: certified seed; host resistance; roguing; selection of seed, cuttings or plants (referred to as seed selection); and management of vectors, pathogens, and alternative hosts (Blomme et al. 2014; Legg 1999; Thomas‐Sharma et al. 2016). Grouping management components by their mode of action can facilitate decision-making and generalization of results. One approach is to group management components into those reducing initial inoculum for each new planting, those reducing the rate of disease spread within a crop, and those reducing the time of exposure of the crop (Berger 1977). Another approach is to group components based on their selectivity against pathogens and whether they manage internal or external inoculum sources (Jones 2006). To compare the performance of management components in this risk assessment framework for seed degeneration, we group components as incidence and rate-altering management components, a logical distinction based on the structure of our models. Incidence-altering management components affect the availability of inoculum from host material in the field or seed lot. For example, roguing (removal of symptomatic plants from the field) affects disease incidence in the field, while seed selection (selecting asymptomatic/least-symptomatic plants as the seed source each season) and use of certified seed, alter disease incidence in the seed lot. Rate-altering management strategies affect the rate of spread of disease in a field, and include strategies such as use of host resistance and vector or pathogen management. (Management of alternative hosts could be treated as reducing inoculum availability from hosts, or as reducing the rate of spread, depending on the goals of an analysis.)

The importance of seed degeneration and the high cost of multi-year field experiments to support empirical analyses have motivated several studies using analytical and simulation models to better understand the process of seed degeneration. Several of these studies have focused on management strategies for cassava diseases, and vector dynamics (Fargette and Vie1995; Holt et al. 1997), illustrating how small management improvements can reduce the risk that a cropping system approaches a threshold for rapid disease increase. In cassava mosaic disease (CMD), weather conditions can affect symptom expression (Gibson and OtimNape 1997), so symptom-based management like roguing and seed selection can have variable success rates from one season to the next. The level of farmer skill in identifying symptoms, especially for pathogens that produce cryptic foliar symptoms, e.g., cassava brown streak disease (CBSD), is another source of variation from one field to another. McQuaid et al. (2016) evaluated the likely performance of roguing for CBSD in seed multiplication sites, showing the potential in sites with low disease pressure. In a more general modeling study, van den Bosch et al. (2007) found that management components such as *in vitro* propagation, high accuracy cutting selection, and use of tolerant varieties, can inadvertently select for virus strains that build-up a high titer in host plants. New analysis of how strategic integration of management components enhances management performance can build on these studies.

Weather is another factor determining the rate of seed degeneration. Viral degeneration of seed potato is lower at high altitudes (Rahman and Akanda 2008), due at least in part to lower virus and vector activity (Fankhauser 2000), and higher rates of reversion or efficiency of autoinfection (Bertschinger 1992). In a fine-scale forecasting model of potato viruses, Bertschinger et al. (1995) used daily temperature measurements to determine host growth rates and vector dynamics, predicting the number of infected progeny seed. In most other models of seed degeneration, however, weather is implicitly addressed in vector dynamics, and weather variability is rarely considered. Understanding the effect of season-to-season weather variability is important for evaluating seed degeneration risk, and understanding climate change scenarios, e.g., where increased population growth of potato virus vectors is predicted for summer crops in parts of South Africa (van der Waals et al. 2013).

There are many potential goals for model development, such as providing a good approximation to reality, precise predictions, or general insights into a phenomenon. Typically models will compromise one of these objectives in the pursuit of others (Gross 2013; Levins 1966). Our goal in developing a risk assessment framework for seed degeneration is to provide a *general* assessment of the performance of different management approaches, as well as a framework that can be adapted to applications for specific pathosystems. We build on the modeling studies of seed degeneration discussed above, with an emphasis on evaluating the effects of both weather variability and variability in management implementation. Thus the benefits of combining management components in an integrated seed health strategy can be explored under different weather scenarios.

The limited data available related to the extent and variability of management component adoption, especially in scenarios where seed degeneration is a problem (e.g., low-income countries), can be a challenge for model parameterization. Studies generally report small-scale, site-specific estimates, so there is little information to guide scaling up consideration to regional or larger extents. In many applications where decisions have to be made despite severe data limitations, such as conservation biology, the use of expert opinion to fill information gaps has gained momentum (Mac Nally 2007; Martin et al. 2005; Yamada et al. 2003). ‘Expert elicitation’ is the systematic collection of the wealth of information integrated into scientists’ opinions through the course of their studies of particular systems (Knol et al. 2010). Use in plant pathology has generally been limited to applications such as the use of expert knowledge for cluster sampling of disease incidence (Hughes and Madden 2002). We explored expert elicitationas a tool to provide the frequency distribution of likely parameter values (such as the level of disease resistance deployed) in India and Africa, along with information about the uncertainty due to lack of knowledge about these systems. Because expert elicitation can provide information about the deployment of a management component across farms in a region, the data it provides can be used to scale up model results to evaluate regional management performance.

We develop here a general risk assessment framework for seed degeneration, designed to inform an integrated seed health strategy for vegetatively-propagated crops (Thomas‐Sharma et al. 2016). The objectives of the study were to (1) build on current theoretical understanding of seed degeneration by including stochasticity of both environmental factors and management components, (2) evaluate scenarios where integrated seed health strategies would be more and less successful, and (3) explore the use of expert elicitation as a method to complement traditional empirical data. We used the framework to ask a set of key questions. (1) Certified seed use is sometimes viewed as a “silver bullet” for managing seed degeneration, yet is unavailable to many farmers. For what scenarios can on-farm management perform as well as certified seed use? And for what scenarios is certified seed use of little value without on-farm management? (2) Given the resource limitations of many farmers in low-income countries, is there an epidemiological basis to choose among within-season management components? Which management components would perform better in the presence or absence of external inoculum? (3) Some methods such as seed selection may present challenges for achieving high levels of effectiveness of implementation, due to cryptic symptoms or lack of farmer experience. Farmers may also choose to plant a mixture of healthy and infected seed when healthy seed is limited and reversion possible (Holt et al. 1997). Can combining management components reduce the minimum effectiveness of implementation required for successful seed degeneration management? (In this study, ‘effectiveness’ is generally used to refer to the effectiveness of implementation of a management component, such as the degree of disease resistance, and differentiated from the effect of management on yield, termed management ‘performance’.) (4) In a development context, the focus may lie not only on the average performance of strategies, but also on the tail of the distribution of performance, the farmers who may be experiencing least benefit from strategies. A stochastic model allows evaluation of the relative performance of strategies across the distribution. How should management be modified to reduce losses at the tail?

## METHODOLOGY

### Overview

#### Purpose

Environmental conditions are important risk-factors for seed degeneration, affecting the build-up of pathogens in a plant, the spread of disease within a field and across large geographic areas, and the usefulness of management practices. A new focus in this framework on the conduciveness of weather for seed degeneration, and variability in weather across cropping seasons, is designed to complement insights from previous seed degeneration models. The effectiveness with which management components are implemented by a farmer can also be variable from one season to the next, depending on the available resources. We address seed degeneration defined as pathogen build-up in seed material, while other physiological factors (such as seed age and physical damage/abnormalities) which can also lower the quality of seed are not considered (Thomas‐Sharma et al. 2016). This risk assessment framework for seed degeneration is designed to be broadly applicable to vegetatively-propagated crops/pathosystems and to capture the key seasonal dynamics of seed degeneration (Fig. 1). While this is not anagent-based model, we generally followed a model description format recommended for agent-based models (Grimm et al. 2010), to enhance clarity and reproducibility. An interactive interface, built by Y. Xing and S. Thomas-Sharma using the Shiny package in R, allows users to experiment directly with the models described here, by accessing the code used in this analysis. It is available at https://yanru-xing.shinyapps.io/SDAppvX1/.

**Fig.1.**
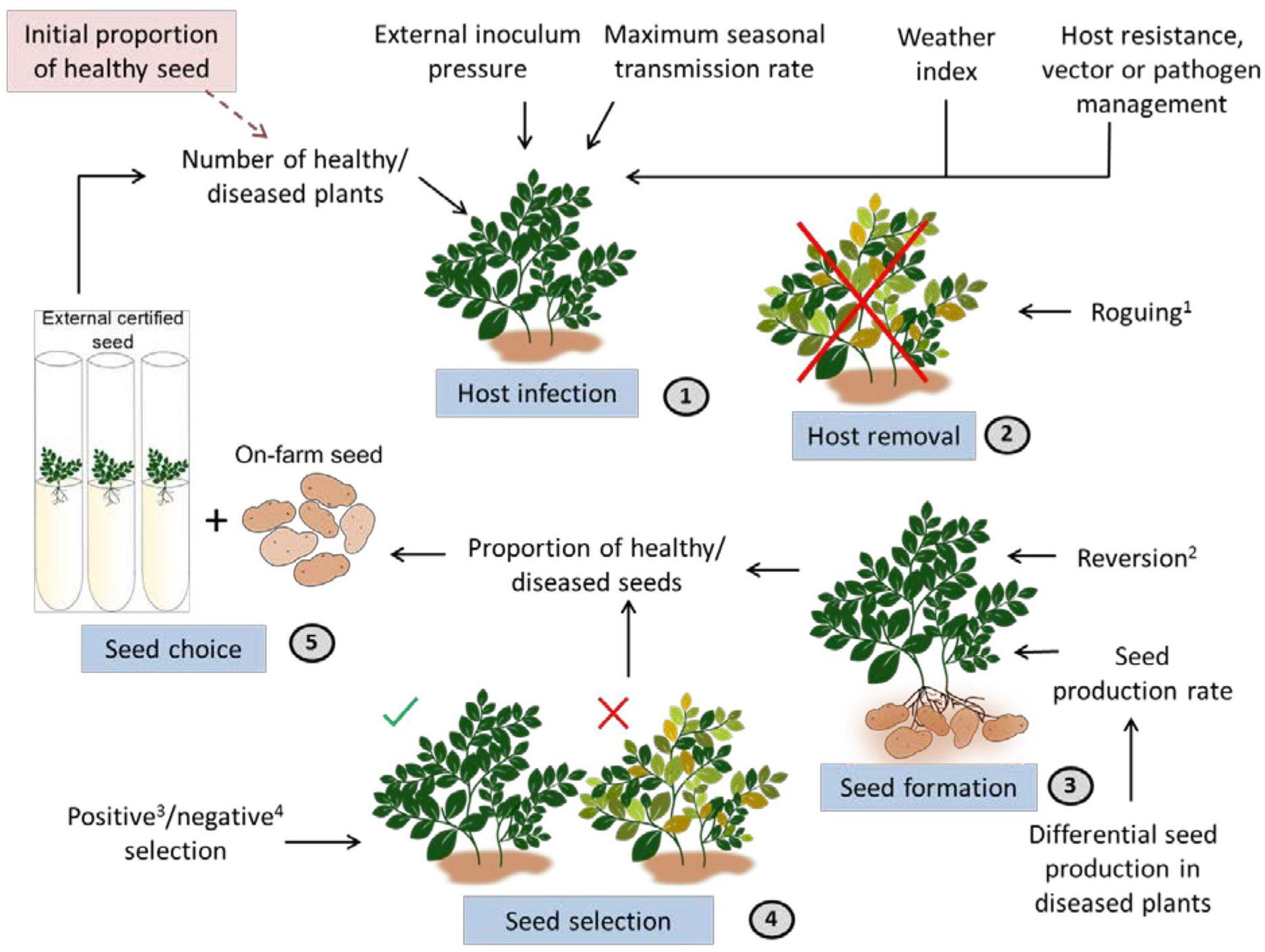
Processes modeled in the seed degeneration risk assessment framework (with potato as an example) are host infection (disease transmission), host removal, seed formation, seed selection, and seed choice. Rate-altering management components (host resistance, vector or pathogen management), incidence-altering management components (seed selection, certified seed usage, and roguing), and phenomena such as reversion and differential seed production in diseased plants modify these processes. The rate of disease transmission is determined by disease-conducive weather conditions (included as a ‘weather index’), external inoculum present, the maximum seasonal transmission rate and any rate-altering management components. In the first season (time-step), the initial proportion of healthy seed used is provided, after which other processes and phenomena are introduced in the order depicted by the circled numbers.

[^1^Removal of diseased plants from the field]
 [^2^Production of disease-free seed by infected mother plant]
 [^3^Selection of asymptomatic plants for seed under high disease intensity]
 [^4^Rejection of symptomatic plants for seed under low disease intensity]

#### Scales and state variables

The model time-step is a season (**
*s*
**), defined as a ‘vegetative generation’, i.e., the time between planting and seed collection during which management decisions are made. For crop species where seeds are collected on a different time scale than harvest of the food crop (e.g., banana or sweetpotato), the production of seed (e.g., banana suckers and sweetpotato vines) and the production of food (e.g., fruit and storage roots) can be considered separately. Seed degeneration is modeled in an individual field without spatially explicit structure, over multiple seasons. Plant and seed populations are characterized by the number or proportion of healthy and diseased individuals, determining the resulting yield loss each season. The state variables are healthy (**HP_*s*_**) and diseased (**DP_*s*_**) plant numbers, healthy (**HS_*s*_**) and diseased (**DS_*s*_**) seed proportions, end of season yield (**Y_*s*_**) and end of season percent yield loss (**YL_*s*_**) (Table 1).

**Table 1.** State variables monitored in seed degeneration risk assessment framework.

| State Variable | Description |
| --- | --- |
| $HP_s$ | Number of healthy plants in field, end of season |
| $DP_s$ | Number of diseased plants in field, end of season |
| $HS_s$ | Number of healthy seed, produced end of season |
| $DS_s$ | Number of diseased seed, produced end of season |
| $pHS_s$ | Proportion of healthy seed, used in following season |
| $pDS_s$ | Proportion of diseased seed, used in following season |
| $Y_s$ | Absolute units of yield, end of season |
| $YL_s$ | Percentage yield loss, end of season |

#### Process overview and scheduling

The model includes five processes that occur every season: host infection, host removal, seed formation, seed selection, and seed choice (Fig. 1). The effects of the following management strategies are evaluated: use of certified seed, host resistance, roguing, seed selection, and vector or pathogen management. (Management of alternative hosts could be evaluated explicitly by incorporating an additional model component, or implicitly as part of the effects of vector or pathogen management.)

1. Host infection, or disease transmission, increases disease incidence in the field, and is a function of the disease incidence in the seed and the availability of external inoculum. The rate of disease transmission is determined by the maximum seasonal disease transmission, the degree to which weather conditions are disease-conducive, any external inoculum present, and the levels of rate-altering management applied in the field (i.e., host resistance, vector or pathogen management). A subset of analyses highlight the greater impact of early-season infections compared to late-season infections. Good proxies for the level of external inoculum are challenging to obtain; in this framework, we included external inoculum as a factor that acts comparably to the presence of infected plants within the field.
2. Host removal occurs by roguing, where diseased plants are removed from the population (death due to disease is treated as minimal). In a subset of analyses, where early and late-season infections are considered, we also highlight the effects of roguing conducted early versus late in the season. Specifying a minimum yield (*minY*) greater than zero supports analysis of the yield penalty due to roguing (when diseased plants produce usable yield). Any compensatory yield effects when roguing is applied (when surrounding plants compensate for yield loss; Salazar 1996) have not been considered.
3. We use the term ‘seed formation’ to describe the production of seed, where the health of the mother plant determines the health of the seed. During seed formation, reversion causes a proportion of diseased plants to become disease-free, producing healthy seed. Diseased plants may produce less seed, contributing less diseased seed to the total on-farm seed produced.
4. Seed selection is represented by a change in the proportion diseased seed produced as a result of selecting against diseased plants as the seed source each season. We do not explicitly describe a distinction often made in seed selection, between positive selection (selection of asymptomatic plants for seed under high disease intensity) and negative selection (rejection of symptomatic plants for seed under low disease intensity). In this model, the proportion of diseased seed is reduced, which might be due to either positive or negative selection.
5. Seed choice affects the proportion of on-farm seed that is combined with certified seed and used in the next season. The model is not spatially explicit, so the degree of mixing among all seed planted in the field is assumed not to have an important effect.

### Design concepts

#### Stochasticity

Two general components are stochastic in this model: seasonal weather-conduciveness for disease, and the effectiveness with which management strategies (vector or pathogen management, seed selection, and roguing) are implemented. The parameters describing these components are the weather index (***W***), the proportional change in infection rate due to vector or pathogen management (***M***), the proportional selection against diseased seed (***Z***), and the proportion diseased plants remaining after roguing (***A***). Each of these follows a normal distribution truncated between 0 and 1, where realizations below 0 are treated as 0, and realizations above 1 are treated as 1.

For the weather index, the mean represents mean climatic conditions, and variation around the mean represents season-to-season variability in conduciveness to disease. For vector or pathogen (or alternative host) management, the mean represents mean effectiveness with which practices are applied and stochasticity captures season-to-season variability, due to timing of application or incomplete control (e.g., changes in the timing and choice of insecticides for vector management). For management practices based on symptom recognition (roguing and seed selection), the mean indicates the mean effectiveness with which the practice is appliedduring a season. Stochasticity captures both variability in symptom expression (e.g., due to variability in timing of infection among seasons, or delayed symptom development) and variability in farmers’ skill in recognizing symptomatic plants. Thus stochasticity in these analyses generally represents what Oberkampf et al. (2004) refers to as ‘aleatory uncertainty’. Similar analyses could also be interpreted in terms of ‘epistemic uncertainty’ or uncertainty due to lack of system knowledge (Oberkampf et al. 2004), or a combination of these two types of uncertainty.

#### Calibration and rate of disease transmission

We conceptualize ***β*** as the maximum rate of disease transmission during the growing season, associated with a scenario where there are no limiting factors for disease spread (i.e., when there is no vector or pathogen management, a highly susceptible host is planted, and the weather is highly disease-conducive). This rate is not necessarily intuitive, because it is multiplied by the number of diseased and healthy plants, in addition to being modified by parameters reflecting the effects of vector or pathogen management, host resistance, and weather. ***β*** is determined by vector and pathogen attributes and other dispersal characteristics, and is interpreted in the context of this general framework as reflecting the maximum rate in the absence of limiting factors. In most simulation experiments, we took ***β*** = 0.02 as the maximum disease transmission rate per season. After exploring the behavior of ***β*** at high and low starting levels of infection with and without management strategies, ***β*** = 0.02 was selected to provide a range of outcomes for evaluation. Substantially lower or higher values of ***β*** resulted in consistent lack of disease, or immediately high disease levels, respectively. Identifying a value of ***β*** through this type of calibration met the needs of our general analysis. However, when developing a more precise application of this framework formanaging a specific crop, calibrating ***β*** for the pathosystem and relevant environments will be an important step.

#### Observations

Model output for a given parameter combination includes summary statistics (mean, 5^th^ percentile, and 95^th^ percentile) for the state variables, and timing for renewal with certified seed, i.e., the first time point at which the proportion of healthy seed falls below a threshold value, where a threshold proportion of 0.7 was used in examples.

### Details

#### Initialization

The initial proportion of healthy seed (*pHS0*) determines the starting infection level in the field, in the first time-step. A relatively low proportion initial infection (0.2) was assumed in most scenarios, and in some cases was compared with a high proportion infection (0.8). Such high pathogen incidence in planting material is a common scenario in low-income countries where farmers routinely use seed of poor health status (Gildemacher et al. 2009), or fields have high disease incidence, making it difficult to select disease-free planting material (Legg 1999).

#### Input data

The current application of the model does not depend on external weather data. However, for more specific applications, the weather index parameter could be defined as a function of a set of observed weather variables relevant to a particular pathosystem.

#### Submodels

There are four submodels that incorporate the effects of weather and management on the state variables (details in Supplementary material S1). The first submodel determines the number of healthy plants at the end of a season as a function of the proportion of diseased seed at the beginning of the season. The second submodel determines the number of healthy seed produced in that season. The third determines yield (in terms of food production) as a function of the proportion of diseased plants. The fourth determines the proportion healthy seed used in the next season.

### Simulation experiments

Simulation experiments were implemented in the R programming environment (R Core Team 2015). Experiments were designed to evaluate the effect of environment, and individual or combined management strategies on yield loss due to seed degeneration, to address the questions posed in the introduction. Each parameter combination was evaluated in 2000 simulations. The maximum (*maxY*) and minimum (*minY*) attainable yields were set at 100 and 0 units, respectively. Some parameters were evaluated at contrasting levels, while other parameters were set to the default values in Table 2 when their effects were not being evaluated. For example, management practices were set to minimum values (i.e., 0), without stochasticity, unless the impact of different levels of effectiveness of implementation was being evaluated. The default values of parameters were selected such that contrasting outcomes could be evaluated. All results are represented such that 0 indicates lack of a management component and 1 indicates complete effectiveness of implementation. (The effects of roguing, seed selection, vector management, and host resistance are likewise described in the results in terms of the effectiveness of implementation, rather than in terms of their corresponding parameter definitions in the model Table 2). The standard deviation for stochastic variables was set to 0.3 and 0.1 for high and low variability scenarios, respectively. Short (5 season) and long-term (10 season) effects on yield loss were studied.

**Table 2.** Parameters used in seed degeneration risk assessment framework

| Parameter | Description | Biological meaning of values | Default values used |
| --- | --- | --- | --- |
| $pHS_0$ | Initial proportion of healthy seed | 1=no seed infected<br>0=all seed infected | 0.8 (low starting infection scenarios)<br>0.2 (high starting infection scenarios) |
| K | Initial plant population (number) | Population at beginning of season based on planting rate in a small field | 100 |
| E | External inoculum | Amount of host/non-host inoculum surrounding a field | 0 (absence of external inoculum)<br>30 (presence of external inoculum) |
| $\beta$ | Maximum transmission rate per season | Maximum rate of disease transmission during the season when there are no limiting factors for disease spread | 0.02 |
| W | Proportional change in infection due to environment | W=1, maximally conducive environmental conditions<br>W=0, environmental conditions that do not support transmission | 0.8 (highly disease-conductive weather)<br>0.2 (marginally disease-conductive weather) |
| $H^1$ | Proportional change in infection due to host genetic resistance | H=1, highly susceptible<br>H=0, immune | 0 |
| $M^1$ | Proportional change in infection rate due to vector management | M=1, indicates no management<br>M=0, indicates vector or pathogen eradication | 0 |
| $A^1$ | Proportion diseased plants remaining after roguing | A=1, indicates no roguing<br>A=0, indicates all diseased plants removed | 0 |
| G | Seed production rate in healthy plants | Number of seed produced per healthy plant | 4 |
| $Z^1$ | Proportional selection against diseased plants (through positive or negative selection) | $Z=1$, indicates no seed selection<br>$Z<1$, indicates proportional selection against diseased plants<br>$Z=0$, indicates complete selection against diseased plants | 0 |
| $C$ | Indicates differential seed production in the diseased plants as a proportion of seed production in healthy plants | $C=0$, indicates no seed production in diseased plants<br>$C=1$, indicates no difference in seed production between healthy and diseased plants<br>$C<1$, indicates reduced seed production in diseased plants<br>$C>1$, indicates increased seed production in diseased plants | 0.9 |
| $R$ | Reversion rate | Proportion of diseased plants that produce disease-free seed | 0.1 |
| $\Phi$ | Proportion certified (or otherwise completely disease-free) seed purchased | $\phi=1$, all certified seed<br>$\phi=0$, no certified seed | 0 |
| $\Theta$ | Rate of decline of end of season yield with increasing disease incidence | $0<\theta\leq 0.5$, indicates yield decline slow initially, then increases<br>$\theta=\text{negative}$, indicates yield decline is rapid initially, then slows<br>$\theta=0$, indicates constant rate of decline | 0.2 |
| $\gamma$ | Proportional change in effect of disease incidence on yield loss for late season versus early season | $\gamma=0$, indicates no yield loss due to late season disease incidence<br>$\gamma=1$, indicates no difference between early and late season effects of disease incidence on yield loss | Not used in general models |
| minY | Minimum yield | Units of yield produced by a severely infected plant | 0 |
| maxY | Maximum yield | Units of yield produced by a healthy plant | 100 |

<sup>1</sup> When addressing results, we describe and discuss these management effects in terms of the
effectiveness of management implementation, so that all types of management can be considered
with 1 indicating complete effectiveness of implementation and 0 indicating complete ineffectiveness. In contrast, for H, M, A, and Z, the model and code are constructed such that 1 indicates no limiting factor for infection processes.

### Parameterization based on expert elicitation

The risk assessment framework described to this point is designed to evaluate risk at a particular field, given the environment and management decisions implemented. Expert elicitation was used to assess the adoption rates for individual management components by farmers in a region, as a first step toward scaling up individual farm risk assessments. In total, twenty-five experts(across crops and geographical regions) provided estimates of the frequency with which different management components were implemented with a particular level of effectiveness. For example, experts estimated the field acreage in each of 10 disease resistance categories in regions of Africa and India. The seed degeneration model described above was used to evaluate outcomes for an individual field, providing the frequency of potential outcomes for a given scenario defined by a set of parameter values. To supplement individual field evaluation, the expert elicitation data provide estimates of the frequency with which different scenarios occur. The data from expert elicitation were used to *partially* calibrate the frequency distribution of yield loss in the risk assessment framework for seed degeneration. Expert elicitation provided relatively high confidence information about the frequency with which farmers used particular management techniques, but did not provide high confidence estimates related to underlying transmission rates (because of the inherent difficulty in estimating transmission rates from personal observations). Thus, expert elicitation made this general analysis relatively more realistic, by indicating how likely different scenarios were to occur, but did not provide a precise estimate of yield outcomes. The details of the methods employed in expert elicitation are in Supplementary material S2.

## RESULTS

### Effect of weather on long-term yield loss

The effect of disease-conducive weather conditions on long-term yield loss was first illustrated in the absence of management, and external inoculum, with other parameters set to default. As expected, highly disease-conducive weather causes yield loss to rise quickly, while, under marginally disease-conducive weather, it rises relatively more slowly and has the potential tostay at an acceptable level (Fig. 2). Season-to-season variability in weather causes seasonal fluctuations in yield loss. Under marginally disease-conducive weather, this variability can cause long-term yield reductions to be very high and comparable to that in highly disease-conducive weather conditions.

**Fig. 2.**
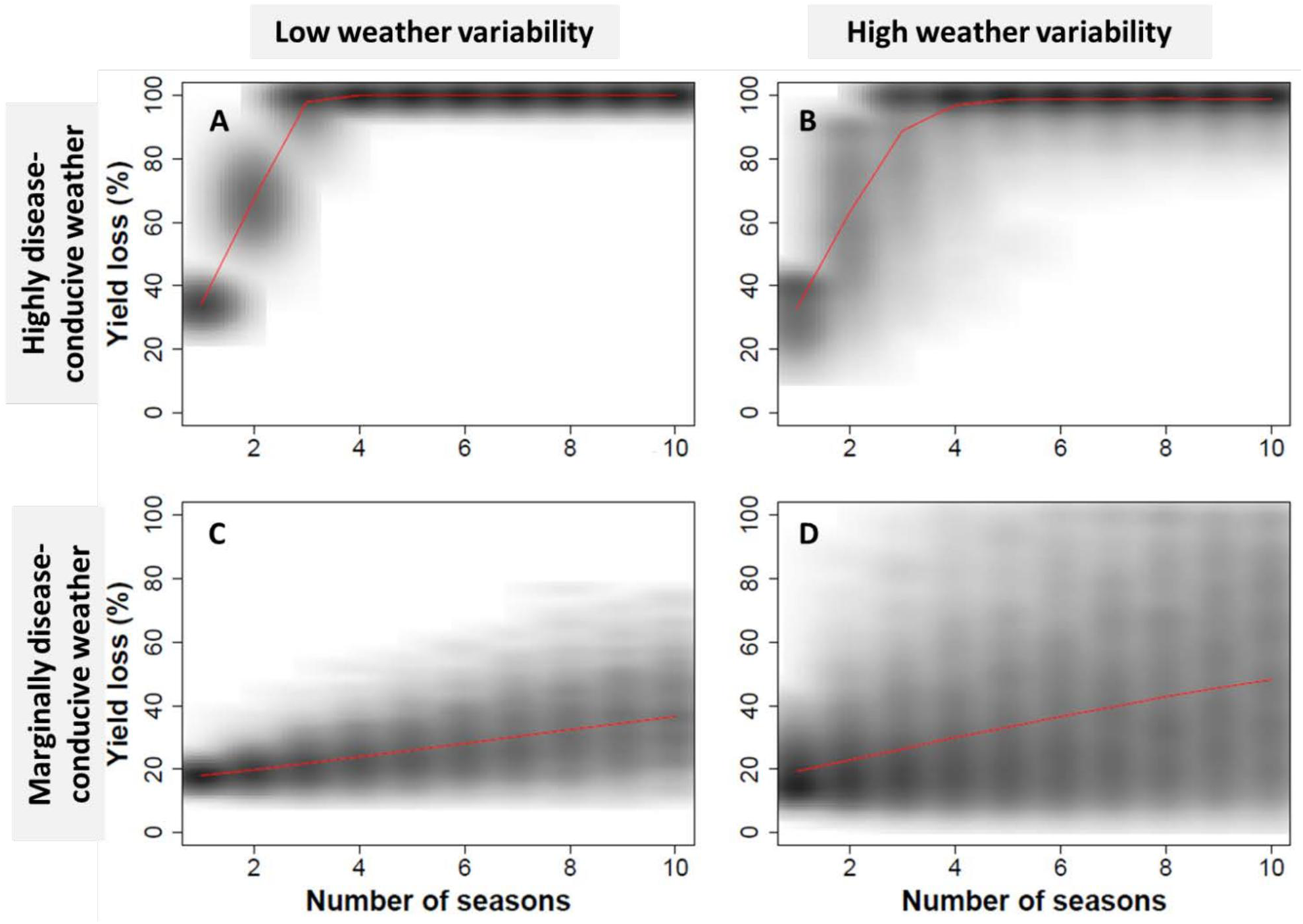
Long-term (10 season) yield loss under no-management scenarios, with low starting levels of infection, under highly and marginally disease-conducive weather scenarios with high (0.3) and low (0.1) season-to-season variability in weather, in the absence of external inoculum (based on 2000 simulations). Other parameters are set to default values from Table 2. Red lines indicate the mean value.

### Effect of individual management practices on yield loss

The effect of individual management practices on short-term yield loss varies with the degree to which weather is disease-conducive (Fig. 3). As disease conduciveness increases, management practices provide less reliable yield loss reduction. For all cases illustrated, under highly disease-conducive conditions, yield loss reaches nearly 100% when the proportional effectiveness of implementation of management practices is low (0-0.2). The effects of the incidence-altering management practices such as roguing, seed selection, and certified seed use are similar to each other. As expected based on the model structure, rate-altering management strategies, such as vector or pathogen management and host resistance, had the same outcome for a given effectiveness of implementation (*not shown separately*).

**Fig. 3.**
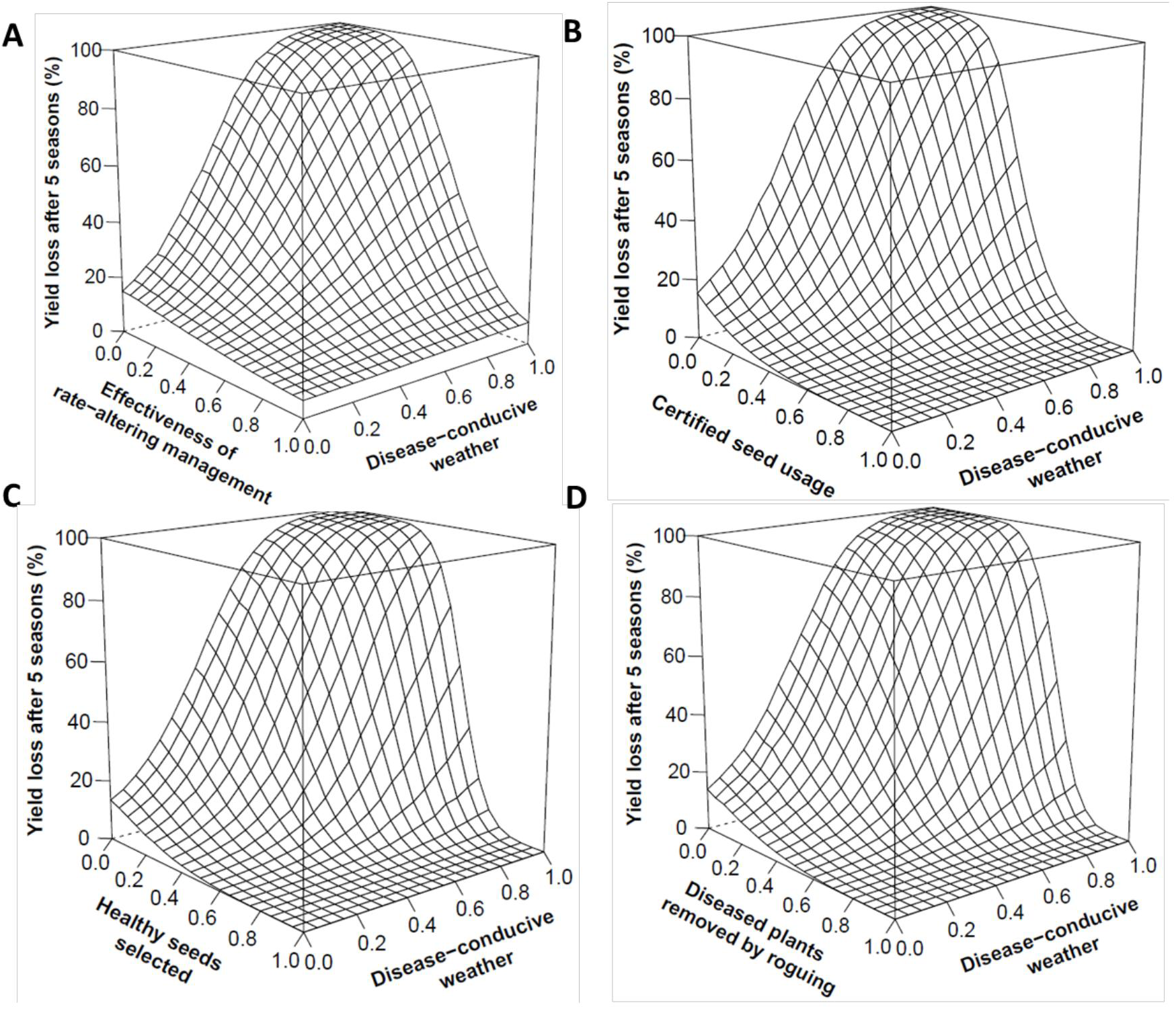
Effect of rate-altering management components (such as vector or pathogen management and host resistance) (A) and incidence-altering management components–certified seed usage (B), seed selection (C), and roguing (D) on percent yield loss after 5 seasons under varying disease-conducive weather conditions, high variability in weather (0.3) and low starting levels of infection, in the absence of external inoculum (based on 2000 simulations). Vector management, seed selection, and roguing assume low variability in effectiveness and are expressed as the proportion effectiveness of management implementation (1 indicating complete effectiveness, and 0 indicating no management). Other parameters are default values from Table 2.

In the absence of external inoculum, strategies such as roguing, use of certified seed and seed selection could substantially reduce yield loss when implemented at 0.2-0.4 proportional effectiveness, under marginally disease-conducive conditions (Fig. 3). Rate-altering management strategies, however, required higher levels of proportional effectiveness of implementation (0.4-0.6) to provide a comparable effect on yield loss. Even when rate-altering management strategies were implemented at ‘complete’ proportional effectiveness (i.e., at 1), in marginally disease-conducive weather conditions, a low level of yield loss (~10%) was observed (Fig. 3A). This was because it took more than 5 seasons for rate-altering management to reduce yield loss levels to zero (*data not shown*). Depending on weather conduciveness and resistance levels, managementpractices such as roguing, use of certified seed and seed selection were thus 20-40% more beneficial than rate-altering management strategies, in the absence of external inoculum (Table 3).

**Table 3.** Minimum effectiveness of management practices required to keep average yield loss below 10% after 10 seasons, under different combinations of weather conduciveness, host resistance, and low starting levels of infection, in the absence of external inoculum (based on 2000 simulations). Host resistance is expressed as 0 for the highly susceptible host and 0.6 for a moderately resistant host. For each of the other management components tested, a range of values from 0 to 1 at increments of 0.1 was evaluated. Other parameters had default values from Table 2.

| Management practice | Highly disease-conducive weather |  | Marginally disease-conducive weather |  | Interpretation |
| --- | --- | --- | --- | --- | --- |
|  | Susceptible host | Resistant host | Susceptible host | Resistant host |  |
| Vector or pathogen management | 0.9 | 0.7 | 0.7 | 0.0 | Proportional effectiveness of vector or pathogen management |
| Roguing | 0.7 | 0.4 | 0.3 | 0.0 | Proportion diseased plants removed |
| Seed selection | 0.7 | 0.4 | 0.3 | 0.0 | Proportion healthy seed selected |
| Use of certified seed | 0.6 | 0.3 | 0.3 | 0.1 | Proportion certified seed used for planting |

When external inoculum is present, however, incidence-altering management was less successful than rate-altering management strategies, reversing the ranking observed in the absence of external inoculum (Fig. 4A, B). When both seed selection and vector or pathogen management were implemented at 0.6 proportional effectiveness, the use of vector or pathogen management in the presence of external inoculum (Fig. 4D) resulted in a relatively slower increase in long-term yield loss compared to seed selection (Fig 4C).

**Fig. 4.**
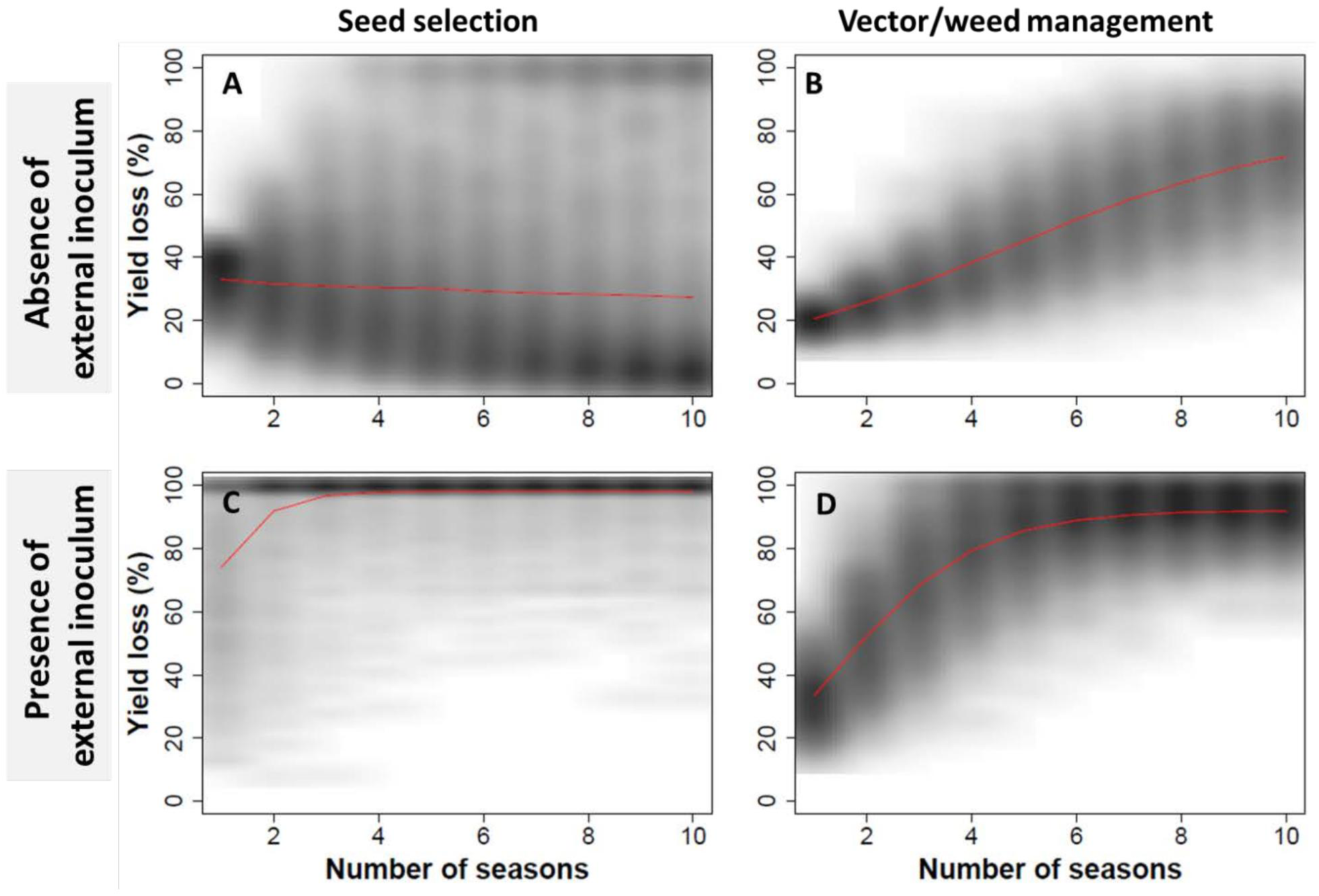
Long-term (10 season) yield loss under seed selection and vector or pathogen management, in the presence and absence of external inoculum (based on 2000 simulations). Vector management and seed selection, expressed in terms of the proportion effectiveness of implementation, had low variability (0.1) and were each set to 0.6 effectiveness of implementation. Other parameters are default values from Table 2. Red line indicates the mean value.

### Effect of combining management strategies on yield loss

The minimum level of effectiveness of implementation for a management component to keep long-term yield loss below 10% (in the absence of external inoculum), changed with the level of resistance used (Table 3). Under highly disease-conducive weather conditions, when susceptible varieties were grown, vector or pathogen management, roguing, seed selection and external certified seed had to be used at 0.9, 0.7, 0.7 and 0.6 proportional effectiveness, respectively, to maintain yield loss <10%. If a resistant variety was used, however, this minimum effectiveness of implementation could be lowered (Table 3). In scenarios where starting infection is high and weather is highly conducive for disease, seed selection is insufficient to keep yield loss below 10% in susceptible varieties (*data not shown*).

Combining management strategies is also useful to delay the need for seed renewal from off-farm certified sources (Table 4). Consider a scenario where renewing seed material with off-farm certified seed becomes necessary when the healthy seed proportion falls below a threshold of 0.7 (which corresponds to approximately 30-40% yield loss depending on conduciveness of weather). In the presence of external inoculum and highly disease-conducive weather conditions, seed renewal was necessary every season when seed selection and vector or pathogen management were practiced individually, but when these practices were combined seed renewal was not necessary for ~12 seasons. In this case, there was strong synergy in the sense that the time to seed renewal for the combined management was substantially larger than the sum of the times to renewal for the two components individually.

**Table 4.** Mean number of seasons until proportion of healthy seed falls below 0.7, in the absence of external inoculum, when a maximum of 15 seasons are considered (based on 2000 simulations). Host resistance and seed selection were evaluated at 0 or 0.6 proportional effectiveness of management. Other parameters had the default values from Table 2.

| Proportional effectiveness of management |  | Highly disease-conducive weather |  | Marginally disease-conducive weather |  |
| --- | --- | --- | --- | --- | --- |
| Seed selection | Host resistance | Absence of external inoculum | Presence of external inoculum | Absence of external inoculum | Presence of external inoculum |
| 0 | 0 | 1.2 | 1.0 | 6.1 | 2.1 |
| 0.6 | 0 | 11.6 | 1.2 | >15.0 | 8.6 |
| 0 | 0.6 | 3.1 | 1.2 | 14.9 | 5.6 |
| 0.6 | 0.6 | >15.0 | 11.6 | >15.0 | >15.0 |

### Effect of season-to-season variability in weather and management practices

Under high proportional effectiveness of implementation (>0.8), high season-to-season variability in vector or pathogen management (*data not shown*) or seed selection resulted in greater yield loss under highly disease conducive weather conditions (Fig. 5). In marginally disease-conducive weather (<0.2) and low proportional effectiveness of implementing seed selection (<0.2), high variability in selection (Fig. 5 B, D) resulted in lower yield loss than low variability scenarios (Fig. 5 C, D). This was because, given the model structure, at low effectiveness of implementation, variability resulted in a higher proportion of healthy plants being incorporated. Conversely, under high effectiveness of implementation, variability in selection resulted in the incorporation of more diseased plants. These trends were more predominant when the starting infection-levels were high (Fig. 5 C, D).

**Fig. 5.**
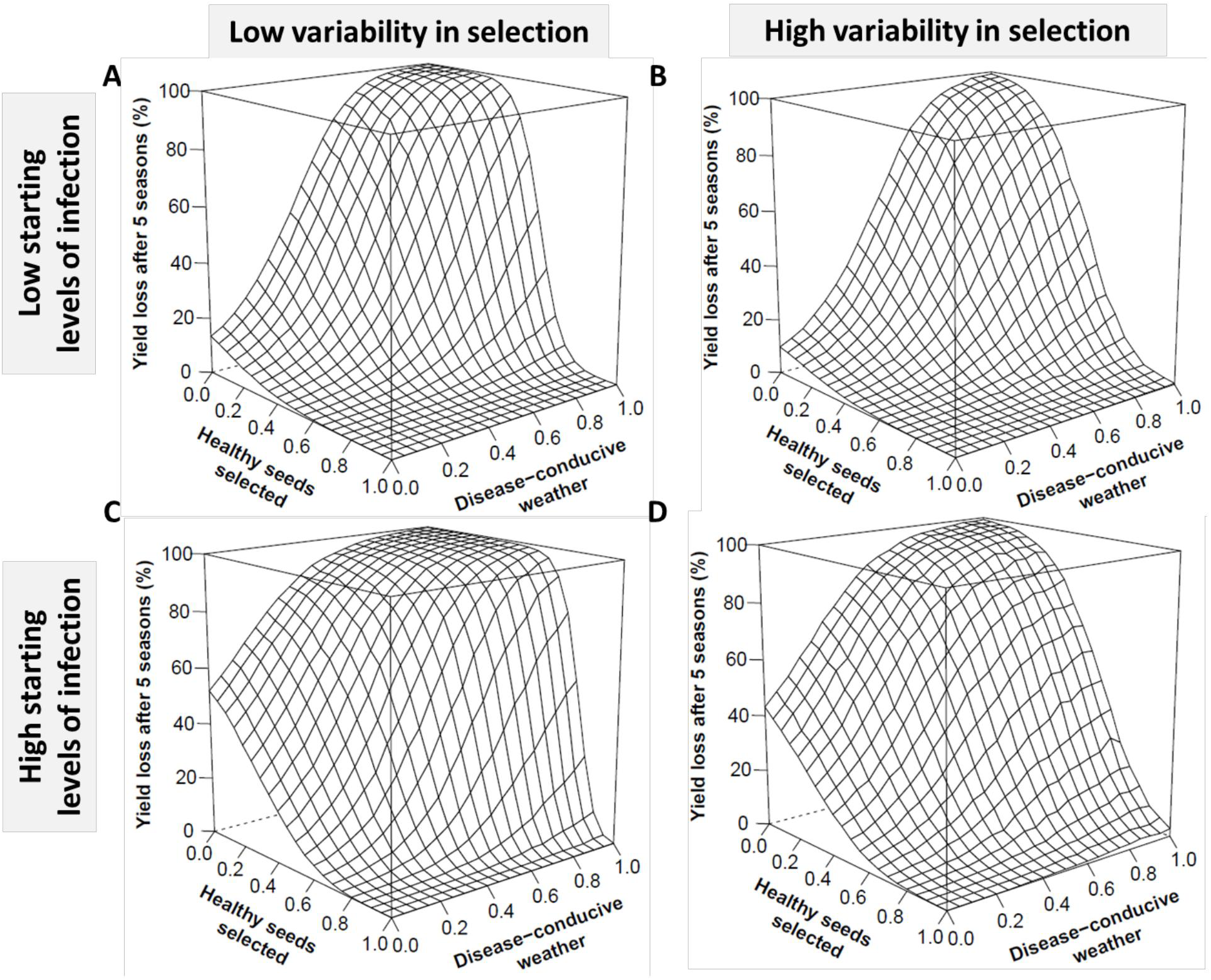
Percent yield loss after 5 seasons as a function of the mean effectiveness of seed selection (proportion healthy seeds selected) for the range of potential levels of disease-conduciveness of weather, at low (A, B) and high (C, D) starting levels of infection, and low (0.1; A, C) and high (0.3; B, D) variability in selection, in the absence of external inoculum (based on 2000 simulations). Seed selection is expressed in terms of the effectiveness of implementation. Other parameters are default values from Table 2.

Season-to-season variability in weather and management practices resulted in variable levels of yield loss (Table 5). In addition to the mean outcomes, we considered the near worst-case outcomes (5th percentile) and the near best-case outcomes (95th percentile). In the near best-case outcome, by implementing seed selection at 0.6 proportional effectiveness for a variety with resistance at level 0.6 out of 1.0, a farmer incurred a yield loss of 16% under highly disease-conducive weather conditions, in the presence of external inoculum (Table 5). However, in theworst-case outcome, implementing management components at the same level of effectiveness resulted in 50% yield loss (Table 5). In the absence of external inoculum, combining seed selection and host resistance resulted in <5% yield loss in best-, worst-case and average outcomes (Table 5).

**Table 5.** Yield loss incurred for average, near worst-case, and near best-case outcomes when seed selection and host resistance are used at 0.6 effectiveness of implementation, under highly disease-conducive weather conditions (based on 2000 simulations). Host resistance and seed selection are expressed as the proportion effectiveness of management implementation (1 indicating complete effectiveness, and 0 indicating no management). Other parameters had the default values in Table 2.

| Proportional effectiveness of management |  | Absence of external inoculum |  |  | Presence of external inoculum |  |  |
| --- | --- | --- | --- | --- | --- | --- | --- |
| Seed selection | Host resistance | 5<sup>th</sup> percentile (Near best-case outcome) | Mean outcome | 95<sup>th</sup> percentile (Near worst-case outcome) | 5<sup>th</sup> percentile (Near best-case outcome) | Mean outcome | 95<sup>th</sup> percentile (Near worst-case outcome) |
| 0 | 0 | 89 | 98 | 100 | 95 | 99 | 100 |
| 0.6 | 0 | 5 | 31 | 86 | 84 | 98 | 100 |
| 0 | 0.6 | 25 | 42 | 56 | 65 | 85 | 97 |
| 0.6 | 0.6 | 1 | 2 | 4 | 16 | 31 | 50 |

### Use of expert elicitation to provide input for crop-specific analyses

In the absence of information about geographic deployment of resistance in cassava, each level of resistance might be considered equally likely, as in an uninformative prior in Bayesian analysis. For a uniform distribution of resistance deployment, model predictions for yield loss in a region would be considerably lower than are likely to be observed, given the rarity of resistance deployment reported in expert elicitation. Crop-specific acreage information obtained from experts (Fig. 6A) can be used to estimate regional yield loss. The resulting modified yield loss distribution (Fig. 6C) is one step more realistic for cassava in Africa and India, in this illustration for marginally disease-conducive weather scenarios.

**Fig. 6.**
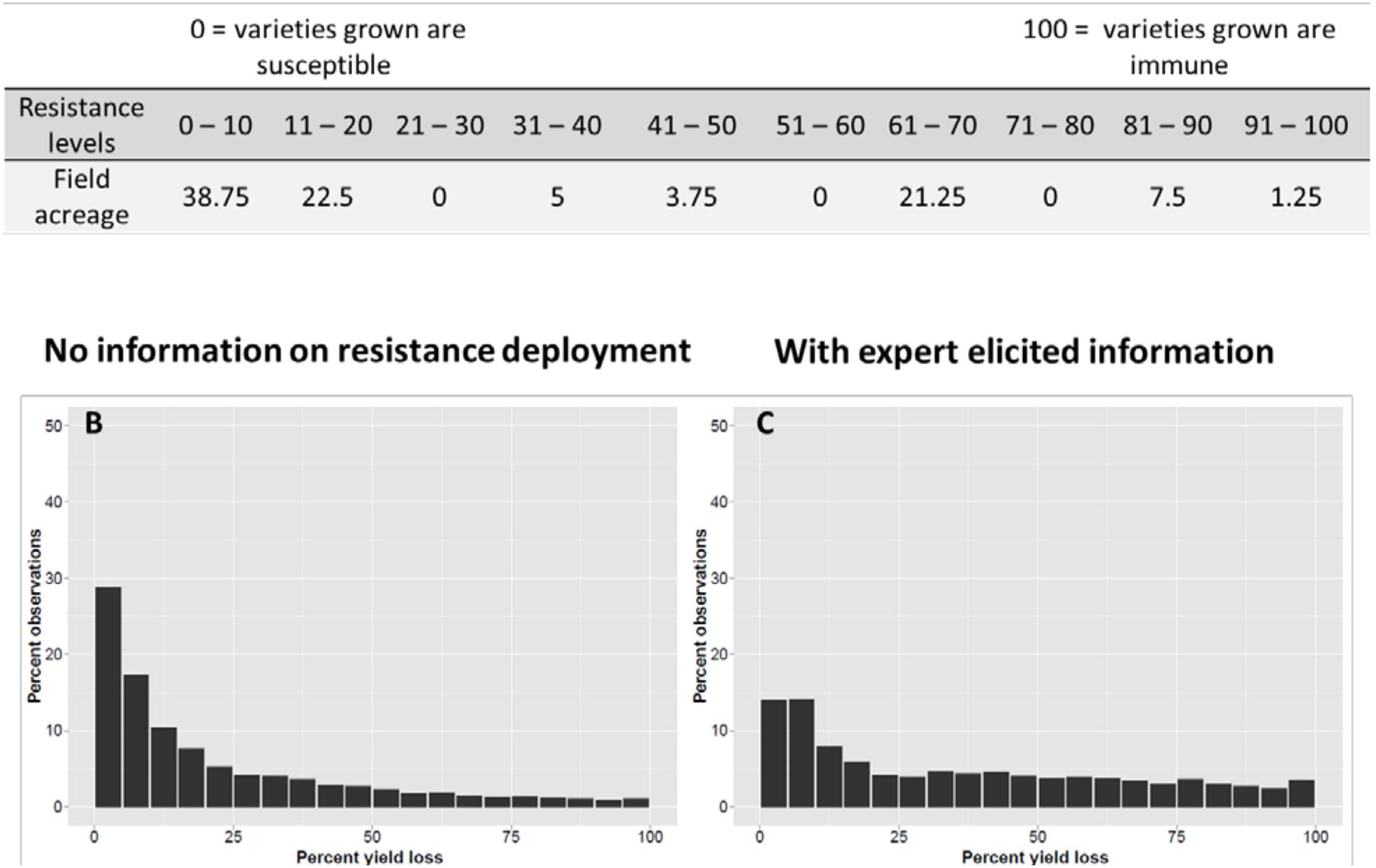
An example of the impact of including expert opinion about the frequency distribution of resistance deployment, compared to an analysis with no information about resistance deployment. A. Mean of experts’ estimate of cassava field acreage under different resistance levels. B. Distribution of yield loss in the absence of information on resistance deployment (i.e., all resistance levels are considered equally likely). C. Distribution of yield loss when experts’ estimate is used to weight resistance levels. The other parameters used in the illustration are pHS0=0.8, K=100, E=0, β=0.02, W=0.3 (high variability [0.3]), M=1 (low variability [0.1]), A=1 (low variability [0.1]), G=4, Z=1 (low variability [0.1]), Y=0.9, R=0.1, φ=0, θ=0.2).

## DISCUSSION

The seed degeneration risk assessment framework was designed to identify scenarios in low-income countries where on-farm management components may be useful, and where they may be absolutely necessary to slow or reverse seed degeneration. We observed that:

*(1) On-farm seed selection can perform as well as certified seed use, if the rate of success in selecting healthy plants is high*. Using the risk assessment framework for seed degeneration, we illustrate how roguing and seed selection can perform as well as use of certified seed (Fig. 3). For many pathosystems, achieving a suitably high rate of success in symptom-recognition is challenging when symptoms are cryptic or variable. If the effectiveness of implementation islow, high yield loss may result despite practicing seed selection (Fig 5D). For cassava seed degeneration caused by CBSD, above-ground symptoms are cryptic (Legg et al. 2011), while for CMD, symptom expression is much more reliable but depends partially on seasonal weather conditions (Gibson and OtimNape 1997). The usefulness of seed selection has been demonstrated for some diseases. For viral diseases and bacterial wilt in potato, farmer-managed trials of seed selection resulted in a ~30% yield increase, with lower disease incidence (Gildemacher et al. 2011; Schulte-Geldermann et al. 2012). Seed selection also increased the tuberous root yields of CMD-susceptible cassava varieties (Mallowa et al. 2006). In parts of western Kenya where CMD is in its post epidemic phase, there is a resurgence of local landraces that are CMD-susceptible, partly because farmers choose the most vigorous plants as seed sources (Mallowa et al. 2006). Farmers may decide against roguing when diseased plants produce usable yield, limiting the practical usefulness of roguing (Legg et al. 2015; Mallowa et al. 2011; Sisterson and Stenger 2013). Under such situations, restricting roguing to early in the season (Supplementary material S3) and coordinating roguing over regional scales (Sisterson and Stenger 2013) can increase the benefits and potentially the incentives for roguing. Additionally, the model treats certified seed material as completely disease-free. Deep-sequencing techniques have revealed that many plant viruses are yet to be described (Kreuze et al. 2009). Thus accurate certification depends on the characterization of a crop virome which can evolve over time and is currently unstudied for crops in many geographical regions. Finally, it is important to remember that, in many low-income countries, farmers have limited or no access to certified seed, and seed selection with even sub-optimum efficiency may provide yield benefits (Holt et al. 1997).

*(2) When choosing among within-season management components, external inoculum can determine the relative usefulness of incidence and rate-altering management.* Management practices such as vector or pathogen management and host resistance can be particularly useful when external inoculum is high and plants are potentially at risk of rapid infection from surrounding fields (Fig. 4). In sweetpotato, the proximity of and level of inoculum in surrounding fields (a function of the level of host resistance in surrounding fields), affected the incidence of sweetpotato viral disease (Aritua et al. 1999). Fargette and Vie (1995) suggested that phytosanitation (by selection of cuttings and roguing) would be more useful in areas with high inoculum levels or susceptible cultivars, because under low inoculum pressure, the use of resistant cultivars with reversion would sufficiently manage the disease. However, where local inoculum levels are very high, plants would get infected quickly, making selection of cuttings and roguing less feasible. In our evaluation, under low external inoculum (e.g., when crop fields are isolated from each other), and when 10% of the cuttings underwent reversion, incidence-altering management was better for managing degeneration than rate-altering management strategies. In post CMD epidemic areas (where there is reduced inoculum), seed selection along with reversion and natural cross protection by mild viruses together allow farmers to cultivate locally preferred, CMD-susceptible varieties (Mallowa et al. 2006).

*(3) For severe disease scenarios, when implementing management components at high levels of effectiveness is difficult, combining management components can produce synergistic benefits and keep seed degeneration below a threshold*. For *Potato leafroll virus* (PLRV) management, the threshold at which vector management becomes necessary may be modified by using resistant varieties (Difonzo et al. 1995). Our study illustrates how the time until renewal with certified seed was needed was prolonged when seed selection and host resistance were applied together (Table 4). In the presence of external inoculum, it is particularly clear that the performance of the combined strategy would be greater than the combination of simple additive effects of individual components, demonstrating potential synergy. We make a simplifying assumption that the time until renewal with certified seed, and the choice of how to integrate management components, depends solely on yield loss. In reality, many socioeconomic factors such as cost and incentives for management, stakeholder preferences, etc., should also be considered to better understand the factors affecting renewal with certified seed, and adoption rates of integrated management practices more broadly in low-income countries (Parsa et al. 2014).

*(4) Combining management components can close the yield gap between average and near worst-case outcomes caused by weather and management heterogeneity*. In the context of development, there may be particular concern for the worst-case outcomes, such as when particularly disease-conducive years may drive vulnerable farmers out of business. High seasonal fluctuations in weather in a geographic region can result in very high yield reductions in the long-term, despite the region being categorized as marginally disease-conducive (Fig 2). The increased frequency of extreme weather is predicted in many climate change scenarios and can lower the performance of disease management practices (Garrett et al. 2013; Jones 2016; Lamichhane et al. 2015). McQuaid et al. (2016) report scenario analyses where, to keep CBSD infection below 10% in seed farms located in areas with lower inoculum pressure, roguing needed to be conducted frequently (weekly or fortnightly intervals) and at a relatively high success rate (70% or higher). Roguing susceptible varieties can however significantly reduce yields compared to a ‘do nothing’ strategy, largely due to the elimination of plant populations (Mallowa et al. 2011). In such cases where farmers are hesitant to remove plants or have difficulty recognizing cryptic symptoms (Legg et al. 2011), infection rates may become too high for roguing or seed selection to be successful. Yield losses can be much higher for near-worst case outcomes than average outcomes, but can be improved by combining management practices (Table 5).

This risk assessment framework for seed degeneration was designed to capture season-to-season dynamics of degeneration, so the model aggregates within-season characteristics of degeneration in most analyses. The model defines degeneration as an increase in incidence of infected seed in the seed lot and does not capture an increase in pathogen load (e.g., virus titer) within a plant. Other specific attributes of vectors such as inoculation efficiency, acquisition efficiency, vector birth/death rates, etc., that are relevant at finer temporal resolution, have been aggregated in the maximum seasonal transmission rate (*β*) and the parameters that modify it. Thus, when calibrating the model for application to a particular crop species in a particular location, developing good estimates of *β* and modifying parameters will be a key step. Calculating the rate of disease transmission when there are no constraints for disease development provides an estimate for *β* (e.g., scenarios where a highly susceptible variety is grown under highly-conducive weather conditions and without management).

Another goal of this risk assessment framework for seed degeneration is flexibility in adapting to the study of different pathogens causing seed degeneration. Although viruses are a major cause of seed degeneration in many crops, in West African yam, the nematodes *Scutellonema bradys*, *Pratylenchus coffeae*, and *Meloidogyne* spp. are major causes of degeneration (Coyne et al. 2010). In bananas grown in Africa, many nematodes (e.g., *Radopholus similis* and *Pratylenchus coffeae*) and bacterial pathogens (e.g., *Xanthomonas campestris* pv. *musacearum*) readily accumulate and spread via planting material (Blomme et al. 2014). In general, *β* may be substantially higher for vector-borne viruses compared to soil-borne fungal pathogens. Also, the weather index directly modifies *β* and may be conceptualized asdirectly affecting nematodes, fungal pathogens, or the dynamics of virus vectors. The potential effectiveness of implementation of management components may vary widely for management of vectors, fungi, bacteria, and nematodes, and can be modified accordingly.

We used expert elicitation to obtain parameter estimates for use in the seed degeneration risk assessment model. Although expert elicitation cannot replace empirical experimentation, we were interested in exploring expert elicitation as a tool to characterize the frequency of different cropping scenarios in a region, which can then be updated as more direct observations become available. A limitation of data from expert elicitation is its subjective nature, potentially influenced by biases. Using data from expert elicitation, we were able to evaluate the relative effects of other management components, taking into account expert estimates of how commonly and with what level of effectiveness the management components were implemented.

This seed degeneration risk assessment framework was designed to answer general questions about the relative performance of management components, alone and in combination, and to provide a platform to answer ‘what-if’ questions for specific scenarios of a crop, pathosystem, and geographic region. For any given pathosystem, implementing the framework can also help to identify key gaps in current knowledge, where parameter estimates are difficult to obtain, that could be the focus of future field studies (Restif et al. 2012). For example, the regional conduciveness of weather to disease could be evaluated based on general observations of regional disease severity, keeping in mind that crop host availability can also be a limiting factor for disease. If good models of weather effects on vector or pathogen dynamics are available, these could be used to evaluate disease-conduciveness in a more flexible way, with more potential to study the effects of weather variability and climate change, and to partition the effects of weather and host abundance. As parameter estimates for a particular pathosystem become available from field studies, the framework can be used to answer questions about the time until renewal with certified seed becomes necessary, and how effectively management components have to be implemented to keep yield loss below a threshold. Ongoing work with the framework is aimed at expanding it to a regional scale in addition to analysis of individual fields, through added information from the literature, new field studies, and expert elicitation. Individual growers act in networks, within which they intentionally and unintentionally exchange information, seed, and pathogens (Garrett 2012; Moslonka-Lefebvre et al. 2011; Shaw and Pautasso 2014). Evaluating the structure of regional networks for the movement of seed may help in targeting where extension and mitigation are most important (Hernandez Nopsa et al. 2015), and may also help to address the challenge of understanding the role of external inoculum in disease risk within a field. Network concepts may also be extended to consider how landscape structures create links among host species (Cox et al. 2013), an important factor for generalist pathogens and vectors. Another important outcome of a regional framework would be regional or larger-extent maps of the likely performance of different seed degeneration management strategies, as an extension of concepts in species distribution mapping (Franklin 2009). Maps of likely management performance can help to inform prioritization by policy makers and extension groups.

## ACKNOWLEDGMENTS

KAG, STS, and GAF developed the framework and model with input from all authors. STS and KAG wrote the R code. STS performed the computational experiments. STS interviewed experts for the expert elicitation component. STS, KAG, and GAF wrote the manuscript with input from all authors. This work was supported by the CGIAR Research Program on Roots, Tubers, and Bananas, the CGIAR Research Program on Climate Change, Agriculture and Food Security, the Kansas Agricultural Experiment Station (Contribution no. ___), a NSF-BMGF BREAD Idea Challenge Award to STS, US NSF Grant EF-0525712 as part of the joint NSF-NIH Ecology of Infectious Disease program, US NSF Grant DEB-0516046, the National Institute for Mathematical and Biological Synthesis (NIMBioS), and the University of Florida. We thank scientists at the following institutions for their participation in the expert elicitation study: Central Tuber Crop Research Institute, Thiruvananthapuram, India; Banana Research Station, Kannara, India; Kerala Agricultural University (KAU), Thrissur, India; KAU, Thiruvananthapuram, India; National Research Center for Banana, Tiruchirapalli, India; International Institute of Tropical Agriculture (IITA), Dar es Salam, Tanzania; IITA Ibadan, Nigeria; University of Queensland, Brisbane, Australia; International Potato Center (CIP), Quito, Ecuador; CIP, Lima, Peru; CIP, Beijing, China; and Indian Agricultural Research Institute, New Delhi, India.

## LITERATURE CITED

Aritua, V., Legg, J. P., Smit, N., and Gibson, R. W. 1999. Effect of local inoculum on the spread of sweet potato virus disease: limited infection of susceptible cultivars following widespread cultivation of a resistant sweet potato cultivar. Plant Pathology 48:655 661.

Berger, R. D. 1977. Application of epidemiological principles to achieve plant disease control. Annual Review of Phytopathology 15:165–183.

Bertschinger, L. 1992. Modelling of potato virus pathosystems by means of quantitative epidemiology. PhD thesis. Swiss Federal Institute of Technology, Zurich, Switzerland.

Bertschinger, L., Keller, E. R., and Gessler, C. 1995. Characterization of the virus x temperature interaction in secondarily infected potato plants using EPIVIT. Phytopathology 85:815–819.

Blomme, G., Jacobsen, K., Ocimati, W., Beed, F., Ntamwira, J., Sivirihauma, C., Ssekiwoko, F., Nakato, V., Kubiriba, J., Tripathi, L., Tinzaara, W., Mbolela, F., Lutete, L., and Karamura, E. 2014. Fine-tuning banana Xanthomonas wilt control options over the past decade in East and Central Africa. European Journal of Plant Pathology 139:265–281.

Carvajal-Yepes, M., Olaya, C., Lozano, I., Cuervo, M., Castano, M., and Cuellar, W. J. 2014. Unraveling complex viral infections in cassava (Manihot esculenta Crantz) from Colombia. Virus Research 186:76–86.

Cox, C. M., Bockus, W. W., Holt, R. D., Fang, L., and Garrett, K. A. 2013. Spatial connectedness of plant species: potential links for apparent competition via plant diseases. Plant Pathology 62:1195–1204.

Coyne, D. L., Claudius-Cole, A. O., Kenyon, L., and Baimey, H. 2010. Differential effect of hot water treatment on whole tubers versus cut setts of yam (Dioscorea spp.). Pest Management Science 66:385–389.

Difonzo, C. D., Ragsdale, D. W., and Radcliffe, E. B. 1995. Potato leafroll virus spread in differentially resistant potato cultivars under varying aphid densities. American Potato Journal 72:119–132.

Fankhauser, C. 2000. Seed-transmitted diseases as constraints for potato production in the tropical highlands of Ecuador. PhD thesis. Swiss Federal Institute of Technology, Zurich, Switzerland.

Fargette, D., and Vie, K. 1995. Simulation of the effects of host resistance, reversion, and cutting selection on incidence of African cassava mosaic virus and yield losses in cassava. Phytopathology 85:370–375.

Fargette, D., Colon, L. T., Bouveau, R., and Fauquet, C. 1996. Components of resistance of cassava to African cassava mosaic virus. European Journal of Plant Pathology 102:645–654.

Franklin, J. 2009. Mapping species distributions: spatial inference and prediction. Cambridge University Press, Cambridge, U.K.

Frost, K. E., Groves, R. L., and Charkowski, A. O. 2013. Integrated control of potato pathogens through seed potato certification and provision of clean seed potatoes. Plant Disease 97:1268–1280.

Garrett, K. A. 2012. Information networks for disease: commonalities in human management networks and within-host signalling networks. European Journal of Plant Pathology 133:75–88.

Garrett, K. A., Dobson, A. D. M., Kroschel, J., Natarajan, B., Orlandini, S., Tonnang, H. E. Z., and Valdivia, C. 2013. The effects of climate variability and the color of weather time series on agricultural diseases and pests, and decision-making for their management. Agricultural and Forest Meteorology 170:216–227.

Gergerich, R. C., Welliver, R. A., Gettys, S., Osterbauer, N. K., Kamenidou, S., Martin, R. R., Golino, D. A., Eastwell, K., Fuchs, M., Vidalakis, G., and Tzanetakis, I. E. 2015. Safeguarding fruit crops in the age of agricultural globalization. Plant Disease 99:176–187.

Gibson, R. W., and OtimNape, G. W. 1997. Factors determining recovery and reversion in mosaic-affected African cassava mosaic virus resistant cassava. Annals of Applied Biology 131:259–271.

Gibson, R. W., and Kreuze, J. F. 2014. Degeneration in sweetpotato due to viruses, virus-cleaned planting material and reversion: a review. Plant Pathology 64:1–15.

Gibson, R. W., Wasswa, P., and Tufan, H. A. 2014. The ability of cultivars of sweetpotato in East Africa to 'revert' from Sweet potato feathery mottle virus infection. Virus Research 186:130–134.

Gildemacher, P. R., Schulte-Geldermann, E., Borus, D., Demo, P., Kinyae, P., Mundia, P., and Struik, P. C. 2011. Seed potato quality improvement through positive selection by smallholder farmers in Kenya. Potato Research 54:253–266.

Gildemacher, P. R., Demo, P., Barker, I., Kaguongo, W., Woldegiorgis, G., Wagoire, W. W., Wakahiu, M., Leeuwis, C., and Struik, P. C. 2009. A description of seed potato systems in Kenya, Uganda and Ethiopia. American Journal of Potato Research 86:373–382.

Grimm, V., Berger, U., DeAngelis, D.L., Polhill, J.G., Giske, J., and Railsback, S.F. 2010. The ODD protocol: a review and first update. Ecological Modelling 221:2760–2768.

Gross, L. J. 2013. Use of computer systems and models. Pages213–220 in: Levin, S.A., ed. Encyclopedia of Biodiversity, Second Edition, Volume 2, pp. 213–220. Academic Press, Waltham, MA.

Hernandez Nopsa, J. F., Daglish, G. J., Hagstrum, D. W., Leslie, J. F., Phillips, T. W., Scoglio, C., Thomas-Sharma, S., Walter, G. H., and Garrett, K. A. 2015. Ecological networks in stored grain: Key postharvest nodes for emerging pests, pathogens, and mycotoxins. BioScience 65:985–1002.

Holt, J., Jeger, M. J., Thresh, J. M., and Otim-Nape, G. W. 1997. An epidemiological model incorporating vector population dynamics applied to African cassava mosaic virus disease. Journal of Applied Ecology 34:793–806.

Hughes, G., and Madden, L. V. 2002. Some methods for eliciting expert knowledge of plant disease epidemics and their application in cluster sampling for disease incidence. Crop Protection 21:203–215.

Jones, R. A. C. 2006. Control of plant virus diseases. Advances in Virus Research 67:205–244.

Jones, R. A. C. 2016. Future scenarios for plant virus pathogens as climate change progresses. Pages 87–147 in: Advances in Virus Research, Vol 95, M. Kielian, K. Maramorosch and T. C. Mettenleiter, eds.

Knol, A. B., Slottje, P. van der, Sluijs, J. P., and Lebret, E. 2010. The use of expert elicitation in environmental health impact assessment: a seven step procedure. Environmental Health 9:19.

Kreuze, J. F., Perez, A., Untiveros, M., Quispe, D., Fuentes, S., Barker, I., and Simon, R. 2009. Complete viral genome sequence and discovery of novel viruses by deep sequencing of small RNAs: A generic method for diagnosis, discovery and sequencing of viruses. Virology 388:1–7.

Lamichhane, J. R., Barzman, M., Booij, K., Boonekamp, P., Desneux, N., Huber, L., Kudsk, P., Langrell, S. R. H., Ratnadass, A., Ricci, P., Sarah, J.-L., and Messean, A. 2015. Robust cropping systems to tackle pests under climate change. A review. Agronomy for Sustainable Development 35:443–459.

Legg, J. 1999. Emergence, spread and strategies for controlling the pandemic of cassava mosaic virus disease in east and central Africa. Crop Protection 18:627–637.

Legg, J. P., Kumar, P. L., Makeshkumar, T., Tripathi, L., Ferguson, M., Kanju, E., Ntawuruhunga, P., and Cuellar, W. 2015. Cassava virus diseases: Biology, epidemiology, and management. Advances in Virus Research 91:85–142.

Legg, J. P., Jeremiah, S. C., Obiero, H. M., Maruthi, M. N., Ndyetabula, I., Okao-Okuja, G., Bouwmeester, H., Bigirimana, S., Tata-Hangy, W., Gashaka, G., Mkamilo, G., Alicai, T., and Kumar, P. L. 2011. Comparing the regional epidemiology of the cassava mosaic and cassava brown streak virus pandemics in Africa. Virus Research 159:161–170.

Levins, R. 1966. The strategy of model building in population biology. American Scientist 54:421–431.

Mac Nally, R. 2007. Consensus weightings of evidence for inferring breeding success in broad-scale bird studies. Austral Ecology 32:479–484.

Madden, L. V., Hughes, G. Van Den, and Bosch, F. 2007. The study of plant disease epidemics. American Phytopathological Society St. Paul, MN.

Mallowa, S. O., Isutsa, D. K., Kamau, A. W., and Legg, J. P. 2011. Effectiveness of phytosanitation in cassava mosaic disease management in a post-epidemic Area of Western Kenya. ARPN Journal of Agricultural and Biological Science 6:8–15.

Mallowa, S. O., Isutsa, D. K., Kamau, A. W., Obonyo, R., and Legg, J. P. 2006. Current characteristics of cassava mosaic disease in postepidemic areas increase the range of possible management options. Annals of Applied Biology 149:137–144.

Martin, T. G., Kuhnert, P. M., Mengersen, K., and Possingham, H. P. 2005. The power of expert opinion in ecological models using Bayesian methods: Impact of grazing on birds. Ecological Applications 15:266–280.

McGuire, S., and Sperling, L. 2016. Seed systems smallholder farmers use. Food Security 8:179–195.

McQuaid, C. F., Sseruwagi, P., Pariyo, A., and van den Bosch, F. 2016. Cassava brown streak disease and the sustainability of a clean seed system. Plant Pathology 65:299–309.

Moslonka-Lefebvre, M., Finley, A., Dorigatti, I., Dehnen-Schmutz, K., Harwood, T., Jeger, M. J., Xu, X., Holdenrieder, O., and Pautasso, M. 2011. Networks in plant epidemiology: From genes to landscapes, countries, and continents. Phytopathology 101:392–403.

Mwangi, J. K., Nyende, A. B., Demo, P., and Matiru, V. N. 2008. Detection of latent infection by Ralstonia solanacearum in potato (Solanum tuberosum) using stems instead of tubers. African Journal of Biotechnology 7:1644–1649.

Oberkampf, W. L., Helton, J. C., Joslyn, C. A., Wojtkiewicz, S. F., and Ferson, S. 2004. Challenge problems: uncertainty in system response given uncertain parameters. Reliability Engineering & System Safety 85:11–19.

Parsa, S., Morse, S., Bonifacio, A., Chancellor, T. C. B., Condori, B., Crespo-Perez, V., Hobbs, S. L. A., Kroschel, J., Ba, M. N., Rebaudo, F., Sherwood, S. G., Vanek, S. J., Faye, E., Herrera, M. A., and Dangles, O. 2014. Obstacles to integrated pest management adoption in developing countries. Proceedings of the National Academy of Sciences, USA 111:3889–3894.

R Core Team. 2015. R: A language and environment for statistical computing. R foundation for statistical computing.

Rahman, M. S., and Akanda, A. M. 2008. Impact of PVY and PLRV on growth and yield of second generation seed potato. Bangladesh Journal of Plant Pathology 24:19–24.

Restif, O., Hayman, D. T. S., Pulliam, J. R. C., Plowright, R. K., George, D. B., Luis, A. D., Cunningham, A. A., Bowen, R. A., Fooks, A. R., O'Shea, T. J., Wood, J. L. N., and Webb, C. T. 2012. Model-guided fieldwork: practical guidelines for multidisciplinary research on wildlife ecological and epidemiological dynamics. Ecology Letters 15:1083–1094.

Salazar, L. F. 1996. Potato viruses and their control. International Potato Center, Lima, Peru.

Schulte-Geldermann, E., Gildemacher, P. R., and Struik, P. C. 2012. Improving seed health and seed performance by positive selection in three Kenyan potato varieties. American Journal of Potato Research 89:429–437.

Shaw, M. W., and Pautasso, M. 2014. Networks and plant disease management: Concepts and applications. Annual Review of Phytopathology 52:477–493.

Sisterson, M. S., and Stenger, D. C. 2013. Roguing with replacement in perennial crops: Conditions for successful disease management. Phytopathology 103:117–128.

Thiele, G. 1999. Informal potato seed systems in the Andes: Why are they important and whatshould‐ we do with them? World Development 27:83–99.

Thomas Sharma, S., Abdurahman, A., Ali, S., Andrade Piedra, J., Bao, S., Charkowski, A., Crook, D., Kadian, M., Kromann, P., and Struik, P. 2016. Seed degeneration in potato: the need for an integrated seed health strategy to mitigate the problem in developing countries. Plant Pathology 65:3–16.

Bosch, F. van den, Jeger, M. J., and Gilligan, C. A. 2007. Disease control and its selection for damaging plant virus strains in vegetatively propagated staple food crops; a theoretical assessment. Proceedings of the Royal Society B-Biological Sciences 274:11–18.

van der Waals, J. E., Kruger, K., Franke, A. C., Haverkort, A. J., and Steyn, J. M. 2013. Climate change and potato production in contrasting South African agro-ecosystems 3. Effects on relative development rates of selected pathogens and pests. Potato Research 56:67–84.

Winkler, R. L. 1996. Uncertainty in probabilistic risk assessment. Reliability Engineering & System Safety 54:127–132.

Yamada, K., Elith, J., McCarthy, M., and Zerger, A. 2003. Eliciting and integrating expert knowledge for wildlife habitat modelling. Ecological Modelling 165:251–264.

